# Lateral Preoptic Area Neurons Signal Cocaine Self-Administration Behaviors

**DOI:** 10.1101/2021.05.29.446307

**Authors:** Kevin R. Coffey, Vaishnavi Venkat, Mark O. West, David J. Barker

## Abstract

The lateral preoptic area is implicated in numerous aspects of substance use disorder. In particular, the lateral preoptic area is highly sensitive to the pharmacological properties of psychomotor stimulants, and its activity promotes drug-seeking in the face of punishment and reinstatement during abstinence. Despite the lateral preoptic areas complicity in substance use disorder, how precisely lateral preoptic area neurons encode the individual components of drug self-administration has not been ascertained. To bridge this gap, we examined how the firing of single lateral preoptic area neurons correlates with three discrete elements of cocaine self-administration: 1) drug-seeking (pre-response), 2) drug-taking (response), and 3) receipt of the cocaine infusion. A significant subset of lateral preoptic area neurons responded to each component with a mix of increases and decreases in firing-rate. A majority of these neurons encoded the operant response with increases in spiking, though responses during the drug-seeking, taking, and reciept windows were highly correlated.

## Introduction

The lateral preoptic area (LPO) is a hypothalamic structure comprised of glutamate and GABA neurons(Barker et al., 2017; Geisler et al., 2007; Jones, 2005) with diverse and wide-ranging connections to limbic and limbic-related structures such as the lateral habenula (LHb) (Barker et al., 2017; Kowski et al., 2008; Mok & Mogenson, 1972; Yetnikoff et al., 2015; Zahm & Root, 2017), the dorsal raphe (DRN) (Peyron et al., 1997; Weissbourd et al., 2014), ventral tegmental area (VTA) (Geisler et al., 2007; Geisler & Zahm, 2006; Yetnikoff et al., 2015), rostromedial tegmental nucleus (RMTg) (Jhou et al., 2009; Yetnikoff et al., 2015), and Locus Coeruleus (LC) (Luppi et al., 1995).

The LPO participates in a wide variety of behaviors from general locomotion(Reichard et al., 2019; Shreve et al., 1989; Shreve & Uretsky, 1991; Subramanian et al., 2018; Zahm et al., 2014) and thirst(Saad et al., 1996), to reward-seeking(Gordon-Fennell et al., 2020) and aversive processing(Barker et al., 2017; Briski & Gillen, 2001). Most notably, the LPO supports strong intracranial self-stimulation(Bushnik et al., 2000; Gallistel et al., 1985; Nakahara et al., 1999) and excitotoxic lesions of the LPO attenuate the rewarding efficacy of medial forebrain bundle self-stimulation(Arvanitogiannis et al., 1996).

A growing body of evidence suggest that the LPO plays a central role in substance use disorder. The LPO and neighboring portions of the substantia innominata participate in the locomotor activation that occurs following the administration of psychomotor stimulants(Swerdlow et al., 1986), and acute amphetamine administration elicits increased c-fos expression in the LPO(Colussi-Mas et al., 2007), especially in LPO efferents to the VTA. Our previous work has shown that LPO neurons are highly sensitive to the pharmacological properties of cocaine, with some neurons tracking ongoing changes in bodily levels of cocaine(Barker et al., 2015).

Perseverative motivated behavior despite conflict is a key trait of substance use disorders(Ettenberg, 2004) that promotes drug relapse(Cooper et al., 2007). Recent evidence has underscored the LPOs role in driving motivated behaviors in the face of conflict(Reichard et al., 2019). Pharmacological stimulation or inhibition of the LPO disrupts the effects of punishment on future cocaine-seeking behavior and stimulating the LPO is capable of driving cocaine reinstatement, likely through excitation of VTA dopamine neurons and inhibition of VTA GABA neurons(Gordon-Fennell et al., 2020).

Here, we seek to determine precisely which components of cocaine self-administration are processed by the LPO, and how those components are encoded. We predict that if LPO neurons participate in cocaine self-administration, they will exhibit firing rate modulations that correlate with particular components of the behavior. To test this hypothesis, we examined how the firing of single LPO neurons correlates with drug-seeking (pre-response), drug-taking (response), and receipt of the cocaine infusion(Coffey et al., 2015; Root et al., 2013). A significant subset of LPO neurons responded to each component with a mix of increases and decreases in spiking. A majority of these neurons encoded the operant response with increased firing, though responses during the drug-seeking, taking, and reciept windows were highly correlated.

## Materials and Methods

All protocols were performed in compliance with the Guide for the Care and Use of Laboratory Animals (NIH, Publication 865–23) and have been approved by the Institutional Animal Care and Use Committee, Rutgers University.

### Subjects and Surgery

Male Long–Evans rats (n = 18, Charles River, Raleigh, NC) were individually housed. All animals were provided ad libitum access to water and sufficient food to bring them to a pre-operative weight of ~330 g. Subjects were anesthetized with sodium pentobarbital (50 mg/kg, i.p.; Abbott Laboratories, North Chicago, IL, USA) and administered atropine methyl nitrate (10 mg/kg, i.p.; Sigma, St. Louis, MO, USA) and penicillin G (75,000 U/0.25 ml, IM). Anesthesia was maintained with sodium pentobarbital (5–10 mg/kg, IP) and ketamine hydrochloride (60 mg/kg, IP; Fort Dodge Laboratories, Fort Dodge, IA, USA). A catheter was implanted into the right jugular vein and exited through a J-shaped stainless steel cannula. An array of either 16 microwires (Microprobes, MD, USA) arranged in two parallel rows of 8 wires with the medial eight wires targeting the LPO or 2 bilateral arrays each with 2 parallel rows of four wires all targeting the LPO were implanted. Both the J-shaped cannula and array headstage were secured to the skull using 5 jewelers’ screws and dental cement. Animals were housed in the self-administration chamber for the remainder of the experiment and were allowed to recover for at least 7 days prior to training. Chambers were located inside a soundproof, ventilated box that was supplied with white noise (75 dB). During all hours other than self-administration sessions, a 200 μL infusion of heparinized-saline was delivered every 25 min by a computer controlled syringe pump, through a leash connected to the animal’s headstage 24/7, to preserve catheter patency. Occasionally, a brief period of anesthesia was induced by an infusion of methohexital sodium (10 mg/kg, IV) in order to check the animal’s patency.

### Self-administration

Daily cocaine self-administration sessions began with insertion of a nonretractable glass lever on the side wall of the Plexiglas chamber and illumination of the house light within the sound attenuating chamber. To initiate the first trial, a stimulus light over the lever was illuminated. One lever press in the presence of the stimulus light immediately extinguished the stimulus light and initiated an intravenous infusion of cocaine (0.24 mg/0.2 ml infusion over 7.5 s) and a 7.5 s tone (3.5 kHz, 70 dB) which co-terminated with the infusion pump. The stimulus light remained off for 40 s during which lever presses had no programmed consequences. Sessions lasted until 50 infusions had been earned or 6 h had elapsed, whichever occurred first.

### Electrophysiological Recordings

All rats were trained for at least 2 weeks prior to electrophysiological recordings (Figure 1A). Neural signals were led from the rat’s head, through a harness, and then to a fluid and electronic swivel (Airflyte Electronics, Bayonne NJ; Plastics One Inc., Roanoke VA). Electrical signals were sampled (50 kHz/wire), digitized, time stamped (0.1 ms resolution), and recorded using SciWorks software (DataWave Technologies, Longmont CO). Self-administration during recording sessions lasted until rats had earned 50 infusions, at which point contingencies ended and the lever was removed. Neural recording sessions began 30 min prior to the start of self-administration and continued for 1 additional hour after the termination of self-administration contingencies (by removal of the glass lever). Each rat received 1-3 neural recording sessions in order to record each of the 15 non-differential probes one time (5 or 15 probes recorded/session, depending on hardware).

**Figure 1.**
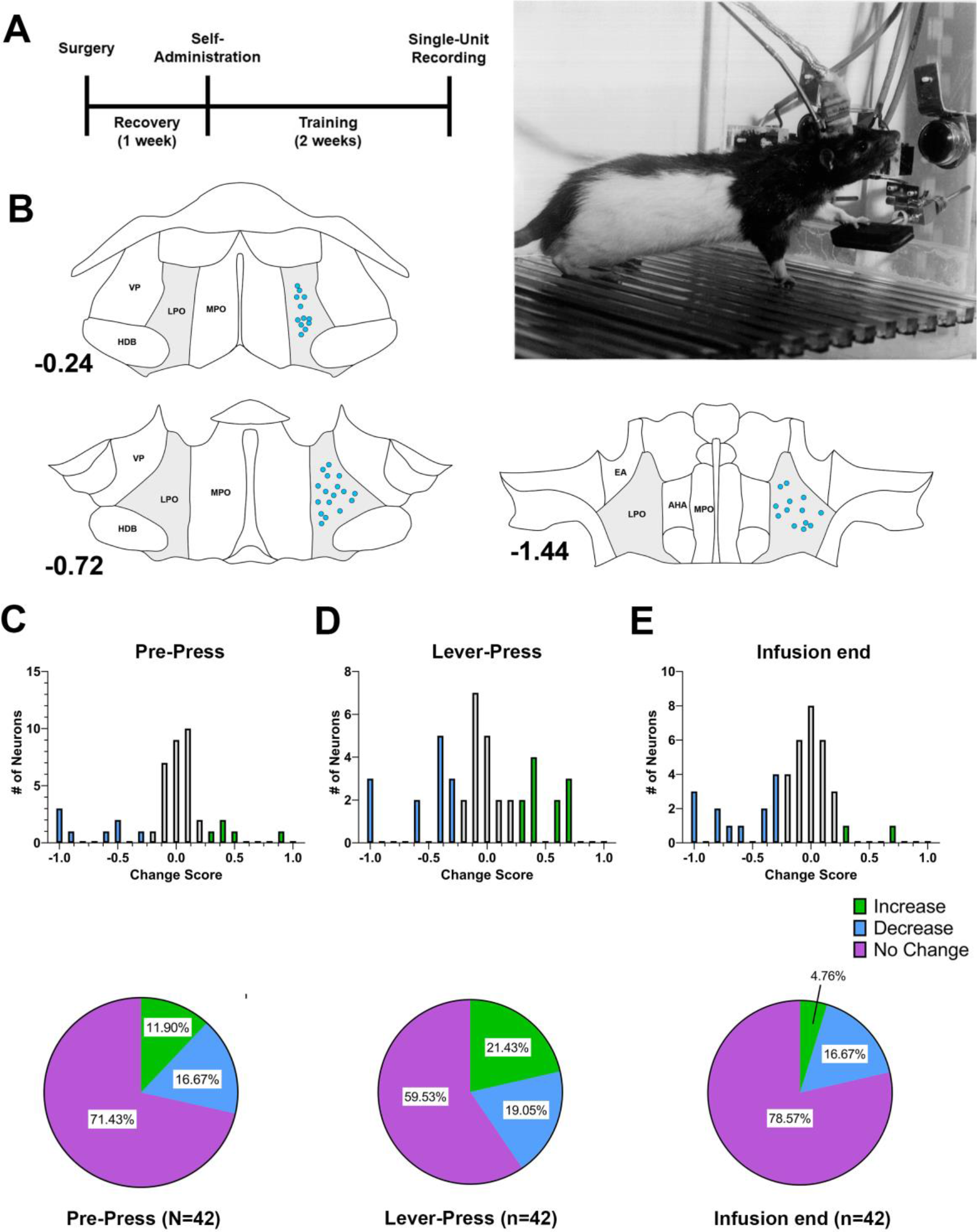
Lateral preoptic area neurons respond to multiple-aspects of cocaine self-administration behavior. **A)** Rats were implanted with an intravenous catheter and then trained to self-administer cocaine for two-weeks prior to electrophysiological recordings of lateral preoptic neurons. **B)** A total of 42 neurons were localized within the borders of the lateral preoptic area. Each blue dot represents the localized tip of a single microwire. **C)** LPO neurons were active during the pre-press window (t (41) = 5.74, p<0.001), with approximately equal numbers of neurons exhibiting pre-press increases (11.90%) or decreases (16.67%). **D)** LPO neurons were most responsive at the time of lever-responding (t (41) = 7.52, p<0.001). ~40% of recorded neurons exhibited responses specific to the lever response. The majority of these were activated at the time of the lever press (21.43%), although other neurons were inhibited at the time of the lever response (19.05%). **E)** The smallest subset of neurons were found to be responsive following the end of the infusion (t (41) = 6.07, p<0.001). These responses were heavily biased towards inhibition (16.67%), with only a small subset of neurons showing post-infusion activation (4.76%).

### Histology

Rats were euthanized with sodium pentobarbital (300 mg/kg, i.p.), and anodal current (50 μA for 4 s) was passed through each electrode. Rats were then perfused with formalin–saline (n = 12) or 4 % paraformaldehyde (n = 6). Brains were post-fixed overnight and then transferred to a 30 % sucrose solution. Sections were taken coronally (50 μm) through the LPO. Tissue was stained in a 5 % potassium ferrocyanide and 10 % HCl and counterstained for antibodies against calbindin D28k immunohistochemistry (ImmunoStar, Inc., Hudson WI), as described previously(Coffey et al., 2015). Wire tracks for every wire in each array were traced from the point at which they penetrated cortex and down to the lesion produced at the microwire tip. In instances where the three-dimensional location of each and every microwire in the array was not able to be accurately tracked, the rat’s data were removed from the dataset. Experimenters blind to results established the location of tracked microwire tips by overlaying microscope images onto a stereotaxic atlas(Paxinos & Watson, 2006) using Photoshop (Adobe, San Jose CA). Wires were included for analysis if they were localized to the LPO. Any microwires localized within 100 μm of any non-LPO brain region were excluded from analysis.

### Isolation/discrimination of individual waveforms

Analysis of neural data was conducted using SciWorks software (DataWave Technologies, Longmont CO). The software was used to isolate neural waveforms from ambient noise and to discriminate different neurons recorded by the same microelectrode. Several measures were taken to identify individual neurons on probes containing more than one waveform. Isolation of neurons was based on multiple measures including spike height, valley and peak amplitudes, valley and peak times, and voltage ranges at selected time points on the ascending and descending limbs of the waveforms. Waveforms were included for analysis if (1) waveforms presented with canonical patterns of neural activity including a clearly defined waveform for the action potential, (2) the amplitude of putative neurons exhibited at least a 2:1 signal/noise ratio, (3) waveform parameters remained stable throughout the entire session, and (4) an interspike interval (ISI) histogram showed that no action potential discharges occurred during the neuron’s natural refractory period (i.e., ~1.4 to 2 ms).

### Statistics

Change scores were calculated for three major event windows: 1) drug-seeking (pre-press) firing patterns, defined as the 500 ms period immediately preceding the lever press, 2) drug-taking (response) firing patterns, defined as the 500 ms period following a lever press, and 3) drug receipt (infusion-related) firing patterns, defined as the 500 ms period immediately following the end of the 7.5s cocaine infusion. Baseline firing was defined as the time window from −10 s to −5 seconds relative to the lever press. Change scores were calculated as:

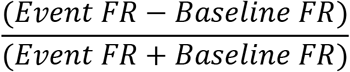

For these event windows, we often observed a bi-directional response of LPO neurons, with a mixture of increases and decreases as compared to each neuron’s baseline firing rate. Thus, we operationally defined neurons exhibiting an ‘increase’ as those showing a two-fold or greater change from baseline firing rates (change score > 0.33) and ‘decreasing’ neurons as those exhibiting a two-fold or greater decrease from baseline firing rates (change score < −0.33). In accounting for the bi-directional nature of neuronal responses, the absolute values of change scores were used for statistical comparisons. Examinations of these neuronal samples were conducted with a one-sample t-test, comparing the population to a value of ‘zero’, which corresponds to a change score representing no change from baseline.

### Results

Out of the 288 total microwires implanted, a total of 42 recorded neurons were localized in the lateral preoptic area (Figure 1B). The average on-drug firing rate of all LPO neurons during baseline was 0.74 ± 0.17 impulses/ second, with the fastest firing neuron reaching 5.49 impulses/second.

### A Subset of LPO Neurons Encode Cocaine-Seeking

The pre-press window encompasses preparatory, drug-seeking behaviors that are proximal to the lever press, including approaches to the lever(Root et al., 2013). A small, but significant number of neurons responded during the pre-press period (t (41) = 5.74, p<0.001; Figure 1 C). The responses of these neurons were heterogeneous, with LPO neurons exhibiting either increases or decreases when compared to their baseline firing rates. In total, 11.90% of LPO neurons exhibited a pre-press increase (n= 5/42), 16.67% of neurons exhibited a decrease (n=7/42) and 71.43% (n=30/42) exhibited no change when compared to baseline.

### A Subset of LPO Neurons Encode Cocaine-Taking

The drug taking period encompasses the moment that rats lever-press for cocaine, along with the presentation of a CS+ cue, and the start of the cocaine infusion. We observed that LPO neurons—as a population—responded at the time of the lever press (t (41) = 7.52, p<0.001; Figure 1D). LPO responses at the time of the lever press were again a mixture of increases or decreases in firing rate when compared to baseline firing rates. More LPO neurons exhibited increases at the time of the lever press (21.43%; n=9/42) as compared to the pre-press drug-seeking period, but a similar number of neurons exhibiting decreases were observed (19.05%; n=8/42). The remaining 59.53% (n=25/42) showed little or no change in their response compared to baseline.

### A Subset of LPO Neurons Encodes Drug Receipt

The drug receipt period encompasses behavioral or neural responses that occur in response to the receipt of the full infusion. The population of LPO neurons was responsive to the receipt of the infusion (t (41) = 6.07, p<0.001; Figure 1E). The primary response following the end of the infusion was an inhibition of LPO neurons, with 16.67% (n=7/42) of LPO neurons exhibiting decreases and only 4.76% (n=2/42) of neurons showing a post-infusion increase. Most neurons exhibited little or no change in their firing rate compared to baseline (78.57%; n=33/42).

These results are largely consistent with the general inhibition of LPO neurons by cocaine over longer time scales(Barker et al., 2015), but also suggest that rapid firing patterns during self-administration behavior can be decoupled from long-lasting changes in neuronal firing associated with the pharmacological effects of cocaine.

### LPO Neuron Activity is Correlated Across Each Component of Cocaine Self-Administration

In examining peri-event responses for LPO neurons across the drug-seeking, drug taking, and drug receipt event windows, we recognized that many neurons exhibited common responses across multiple events. A graphical analysis of neurons inhibited at the time of the lever press revealed that this subset of neurons exhibited a lengthy change in firing rate, in that they were also inhibited at the time of the pre-press and infusion event windows. (Figure 2A). On the other hand, neurons that were excited at the time of the lever press also exhibited excitatory responses during the pre-press window (Figure 2B) but exhibited a more transient change in firing rate and appeared to be unresponsive or mildly inhibited at the end of the infusion.

**Figure 2.**
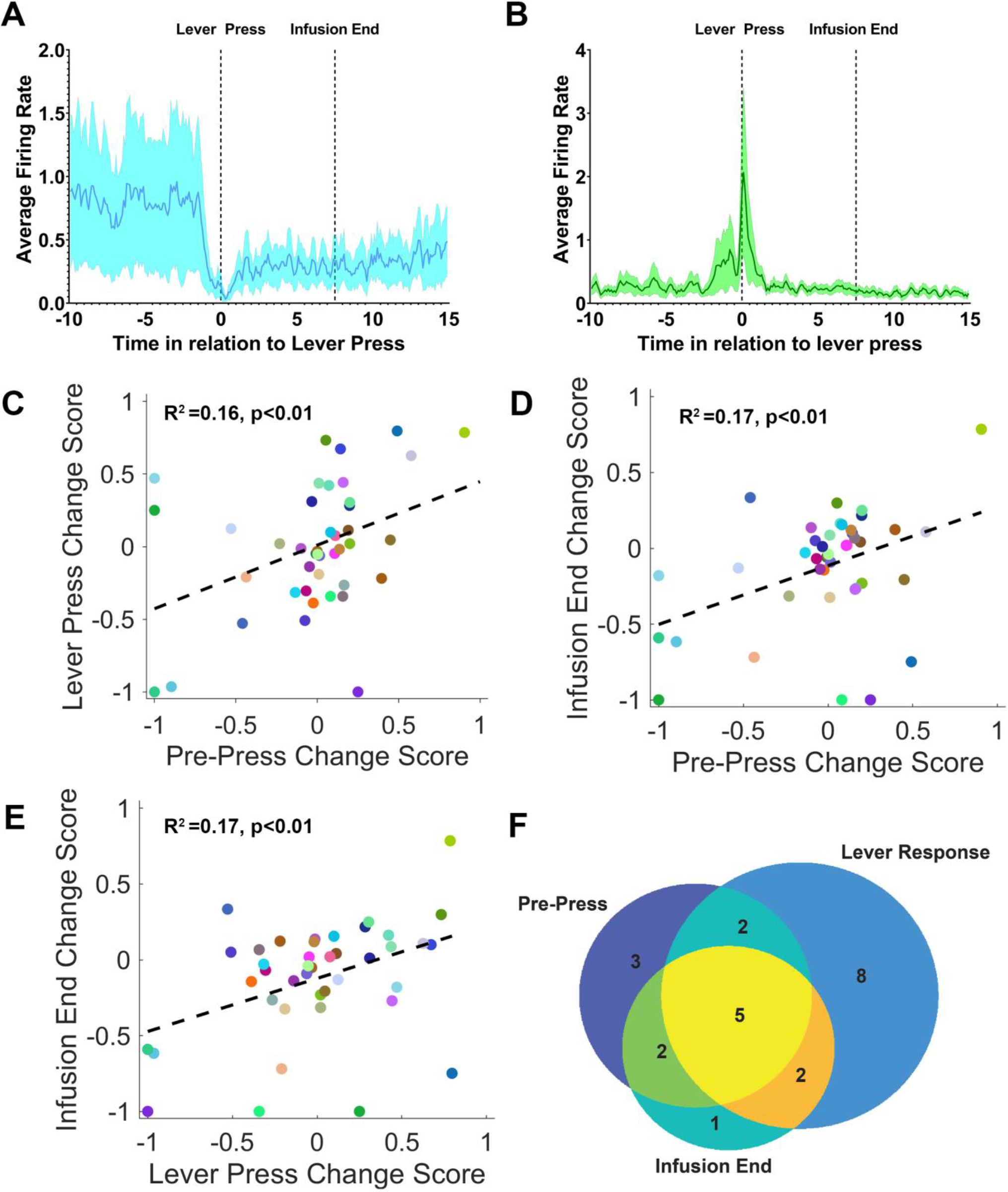
The responses of lateral preoptic area neurons are tied to operant responding. **A)** Averaging neurons exhibiting an inhibitory response at the time of the lever response revealed peri-response inhibition, followed by a long-lasting inhibitory response**. B)** Averaging neurons exhibiting an excitatory response at the time of the lever response revealed peri-response excitation that was short lived, starting during the pre-press window and extending until just after the lever response. **C-E)** Consistent with the observation of common response patterns across multiple events, we observed that responding during the pre-press, lever press, and infusion end windows was highly correlated. Each data point is color-coded for specific neuron of interest to allow for cross-comparison. **F)** In cross-examining responsive, it became clear that responses were most heavily tied to the operant response, with some responses beginning just prior to the operant response and a small number extending across the length of the infusion period.

Accordingly, we observed that responses across events were highly correlated. Responses during the pre-press window were correlated with those at the lever press (R_2_=0.16, P<0.01; Figure 2C), although the range of change scores during the pre-press window was far smaller. Similarly, responses at the time of the pre-press were highly correlated with those at the end of the infusion (R^2^=0.17, p<0.01;). As observed in the average response profile (Figure 2A), this correlation appeared to be driven by long-lasting inhibitory responses. Finally, responses at the lever press were correlated with those at the end of the infusion (R^2^=0.17, p<0.01; Figure 2E). Similar to the relationship observed between the pre-press and lever press, the dynamic range for change scores at the time of the infusion was much smaller.

Based on these findings, we conducted a graphical analysis of neurons that exhibited operationally defined increases, decreases, or a mixture of responses across one or more events (n=23 of 42 neurons). This analysis suggested that most large neuronal changes were tied to the time of the lever responses, with 8 out of the 23 neurons (34.8% of responsive neurons) responding only to the lever press, and an additional 9 neurons (39.1%) also responding during the pre-press window (n=2; 8.7%), infusion window (n=2; 8.7%), or both (n=5; 21.7%). A small subset of neurons responded only during the pre-press window (n=3; 13%) or during both the pre-press and infusion windows (n=2; 8.7%), and only 1 neuron responded only at the time of the infusion window (4.3%). Overall, these results are consistent with the hypothesis that the responses of LPO neurons are intimately tied with drug-taking responses.

## Discussion

The LPO has been repeatedly linked to substance use disorder. Nonetheless, it has never been explicitly demonstrated whether the firing of LPO neurons correlates with drug self-administration (SA) behavior, and to which components of SA these neurons are responsive. Here we provide evidence that subsets of LPO neurons are responsive across multiple events related to operant SA, with most neurons directly encoding drug-taking (lever press) responses. Further, modulation of a small subset of LPO neurons preceded the lever-press, indicating they encode drug-seeking. The responses of LPO neurons were heterogeneous in nature, with neurons that were either excited or inhibited during the pre-press, lever press, and infusion windows. Moreover, we observed that the responses of these neurons are complex, with several neurons exhibiting long-lasting responses that begin prior to the lever press and continue through the end of the infusion.

The presence of two opposing classes of LPO neurons is particularly interesting and may relate to the heterogeneity in LPO neuronal populations. The LPO comprises both glutamatergic and GABAergic neurons(Barker et al., 2017; Geisler et al., 2007; Jones, 2005), but lacks the cholinergic neurons found in adjacent forebrain regions(Gritti et al., 1998). We recently observed that LPO glutamate and GABA neurons play opposing roles in driving motivated behavior, at least in their projections to the lateral habenula, with the activation of GABAergic neurons producing reward and the activation of glutamatergic neurons driving aversion(Barker et al., 2017). Moreover, a recent study observed similar kinds of opposing responses in LPO projections to the ventral tegmental area(Gordon-Fennell et al., 2019). The opposing neuronal subtypes we observed during self-administration may represent a coordinated but opposing response by LPO glutamate and GABA neurons.

Several prior studies have linked the LPO to locomotor responses or drug-induced locomotor responses (Jones & Mogenson, 1980; Mogenson & Nielsen, 1983; Mogenson et al., 1983; Shreve & Uretsky, 1989, 1991). These responses appear to depend on LPO local glutamatergic or GABAergic signaling, with GABAergic and glutamatergic signaling playing opposing roles(Shreve & Uretsky, 1991). Based on these studies, one might conclude that the long-lasting responses observed across the pre-press, lever press, and infusion events could relate to locomotor activity during the peri-response window of self-administration sessions. However, our prior work found no relationship between LPO firing rates and locomotor activity(Barker et al., 2015). Moreover, one recent study failed to find an increase in general locomotor activity in a context associated with operant responding(Gordon-Fennell et al., 2020), and additional work has suggested that this locomotor activity may reflect fixed action patterns(Reichard et al., 2019). When considered alongside the present data, these results suggest that the LPO plays a more explicit role in goal-directed behaviors than was originally thought.

## Conclusion

We provide here the first evidence that LPO neurons participate in multiple aspects of drug self-administration, but most notably drug-taking behavior. Our findings also indicate that only a subset of LPO neurons participate in signaling during cocaine self-administration. It would therefore be of great interest for future studies to examine whether these responsive ensembles comprise a specific cell-type or project to a particular structure; projections to the LHb(Barker et al., 2017), VTA(Gordon-Fennell et al., 2020; Gordon-Fennell et al., 2019), or RMTG(Yetnikoff et al., 2015) are all compelling targets. Additionally, with the recent finding that the LPO may play an important role in drug relapse and decision making during relapse(Gordon-Fennell et al., 2020), more work is needed to determine the roles of specific LPO neuronal subtypes in supporting drug-seeking behavior. A better understanding of these processes could aid in the development of therapies to sustain drug abstinence in the face of repeated conflict.

## Acknowledgements

This study was supported by the National Institute on Drug Abuse Grants DA043572 (DJB) and DA06886 (MOW). The funders had no role in study design, data collection, data analysis, decision to publish, or manuscript preparation. MW and DJB conceptualized the project. DJB and KRC performed the surgeries, electrophysiological recordings, and the histology. DJB, and VV Analyzed the data. KRC and DJB wrote the manuscript with contributions from all co-authors. All authors have seen the manuscript and approved it for publication. The authors declare that they do not have any conflicts of interest (financial or otherwise) related to the data presented in this manuscript.

## Notes

### Competing Interest Statement

The authors have declared no competing interest.

## References

Arvanitogiannis, A., Waraczynski, M., & Shizgal, P. (1996). Effects of excitotoxic lesions of the basal forebrain on MFB self-stimulation. Physiol Behav, 59(4-5), 795–806. doi:10.1016/0031-9384(95)02157-4

Barker, D. J., Miranda-Barrientos, J., Zhang, S., Root, D. H., Wang, H.-L., Liu, B., … Morales, M. J. C. r. (2017). Lateral preoptic control of the lateral habenula through convergent glutamate and GABA transmission. 21(7), 1757–1769.

Barker, D. J., Striano, B. M., Coffey, K. C., Root, D. H., Pawlak, A. P., Kim, O. A., … Function. (2015). Sensitivity to self-administered cocaine within the lateral preoptic–rostral lateral hypothalamic continuum. Brain Structure and Function, 220(3), 1841–1854.

Briski, K., & Gillen, E. J. B. r. b. (2001). Differential distribution of Fos expression within the male rat preoptic area and hypothalamus in response to physical vs. psychological stress. 55(3), 401–408.

Bushnik, T., Bielajew, C., & Konkle, A. T. (2000). The substrate for brain-stimulation reward in the lateral preoptic area. I. Anatomical mapping of its boundaries. Brain Res, 881(2), 103–111. doi:10.1016/s0006-8993(00)02564-6

Coffey, K. R., Barker, D. J., Gayliard, N., Kulik, J. M., Pawlak, A. P., Stamos, J. P., & West, M. O. (2015). Electrophysiological evidence of alterations to the nucleus accumbens and dorsolateral striatum during chronic cocaine self-administration. Eur J Neurosci, 41(12), 1538–1552. doi:10.1111/ejn.12904

Colussi-Mas, J., Geisler, S., Zimmer, L., Zahm, D. S., & Berod, A. (2007). Activation of afferents to the ventral tegmental area in response to acute amphetamine: a double-labelling study. Eur J Neurosci, 26(4), 1011–1025. doi:10.1111/j.1460-9568.2007.05738.x

Cooper, A., Barnea-Ygael, N., Levy, D., Shaham, Y., & Zangen, A. (2007). A conflict rat model of cue-induced relapse to cocaine seeking. Psychopharmacology (Berl), 194(1), 117–125. doi:10.1007/s00213-007-0827-7

Ettenberg, A. (2004). Opponent process properties of self-administered cocaine. Neurosci Biobehav Rev, 27(8), 721–728. doi:10.1016/j.neubiorev.2003.11.009

Gallistel, C. R., Gomita, Y., Yadin, E., & Campbell, K. A. (1985). Forebrain origins and terminations of the medial forebrain bundle metabolically activated by rewarding stimulation or by reward-blocking doses of pimozide. J Neurosci, 5(5), 1246–1261.

Geisler, S., Derst, C., Veh, R. W., & Zahm, D. S. (2007). Glutamatergic afferents of the ventral tegmental area in the rat. J Neurosci, 27(21), 5730–5743. doi:10.1523/JNEUROSCI.0012-07.2007

Geisler, S., & Zahm, D. S. (2006). Neurotensin afferents of the ventral tegmental area in the rat:[1] re-examination of their origins and [2] responses to acute psychostimulant and antipsychotic drug administration. European Journal of Neuroscience, 24(1), 116–134.

Gordon-Fennell, A., Gordon-Fennell, L., Desaivre, S., & Marinelli, M. (2020). The Lateral Preoptic Area and Its Projection to the VTA Regulate VTA Activity and Drive Complex Reward Behaviors. Front Syst Neurosci, 14, 581830. doi:10.3389/fnsys.2020.581830

Gordon-Fennell, A. G., Will, R. G., Ramachandra, V., Gordon-Fennell, L., Dominguez, J. M., Zahm, D. S., & Marinelli, M. (2019). The Lateral Preoptic Area: A Novel Regulator of Reward Seeking and Neuronal Activity in the Ventral Tegmental Area. Front Neurosci, 13, 1433. doi:10.3389/fnins.2019.01433

Gritti, I., Mariotti, M., & Mancia, M. (1998). GABAergic and cholinergic basal forebrain and preoptic-anterior hypothalamic projections to the mediodorsal nucleus of the thalamus in the cat. Neuroscience, 85(1), 149–178. doi:10.1016/s0306-4522(97)00573-3

Jhou, T. C., Geisler, S., Marinelli, M., Degarmo, B. A., & Zahm, D. S. J. J. o. C. N. (2009). The mesopontine rostromedial tegmental nucleus: a structure targeted by the lateral habenula that projects to the ventral tegmental area of Tsai and substantia nigra compacta. 513(6), 566–596.

Jones, B. E. (2005). From waking to sleeping: neuronal and chemical substrates. Trends Pharmacol Sci, 26(11), 578–586. doi:10.1016/j.tips.2005.09.009

Kowski, A. B., Geisler, S., Krauss, M., & Veh, R. W. (2008). Differential projections from subfields in the lateral preoptic area to the lateral habenular complex of the rat. J Comp Neurol, 507(4), 1465–1478. doi:10.1002/cne.21610

Luppi, P. H., Aston-Jones, G., Akaoka, H., Chouvet, G., & Jouvet, M. (1995). Afferent projections to the rat locus coeruleus demonstrated by retrograde and anterograde tracing with cholera-toxin B subunit and Phaseolus vulgaris leucoagglutinin. Neuroscience, 65(1), 119–160. doi:10.1016/0306-4522(94)00481-j

Mok, A. C., & Mogenson, G. J. (1972). An evoked potential study of the projections to the lateral habenular nucleus from the septum and the lateral preoptic area in the rat. Brain Res, 43(2), 343–360. doi:10.1016/0006-8993(72)90392-7

Nakahara, D., Ishida, Y., Nakamura, M., Kuwahara, I., Todaka, K., & Nishimori, T. (1999). Regional differences in desensitization of c-Fos expression following repeated self-stimulation of the medial forebrain bundle in the rat. Neuroscience, 90(3), 1013–1020. doi:10.1016/s0306-4522(98)00510-7

Paxinos, G., & Watson, C. (2006). The rat brain in stereotaxic coordinates: hard cover edition: Elsevier.

Peyron, C., Petit, J.-M., Rampon, C., Jouvet, M., & Luppi, P.-H. J. N. (1997). Forebrain afferents to the rat dorsal raphe nucleus demonstrated by retrograde and anterograde tracing methods. 82(2), 443–468.

Reichard, R. A., Parsley, K. P., Subramanian, S., Stevenson, H. S., Schwartz, Z. M., Sura, T., & Zahm, D. S. (2019). The lateral preoptic area and ventral pallidum embolden behavior. Brain Struct Funct, 224(3), 1245–1265. doi:10.1007/s00429-018-01826-0

Root, D. H., Ma, S., Barker, D. J., Megehee, L., Striano, B. M., Ralston, C. M., … West, M. O. (2013). Differential roles of ventral pallidum subregions during cocaine self-administration behaviors. J Comp Neurol, 521(3), 558–588. doi:10.1002/cne.23191

Saad, W. A., Luiz, A. C., De Arruda Camargo, L. A., Renzi, A., & Manani, J. V. (1996). The lateral preoptic area plays a dual role in the regulation of thirst in the rat. Brain Res Bull, 39(3), 171–176. doi:10.1016/0361-9230(95)02089-6

Shreve, P., Uretsky, N. J. P. B., & Behavior. (1989). AMPA, kainic acid, and N-methyl-D-aspartic acid stimulate locomotor activity after injection into the substantia innominata/lateral preoptic area. 34(1), 101–106.

Shreve, P. E., & Uretsky, N. J. (1991). GABA and glutamate interact in the substantia innominata/lateral preoptic area to modulate locomotor activity. Pharmacol Biochem Behav, 38(2), 385–388. doi:10.1016/0091-3057(91)90296-e

Subramanian, S., Reichard, R. A., Stevenson, H. S., Schwartz, Z. M., Parsley, K. P., & Zahm, D. S. (2018). Lateral preoptic and ventral pallidal roles in locomotion and other movements. Brain Struct Funct, 223(6), 2907–2924. doi:10.1007/s00429-018-1669-2

Swerdlow, N. R., Vaccarino, F. J., Amalric, M., & Koob, G. F. (1986). The neural substrates for the motor-activating properties of psychostimulants: a review of recent findings. Pharmacol Biochem Behav, 25(1), 233–248. doi:10.1016/0091-3057(86)90261-3

Weissbourd, B., Ren, J., DeLoach, K. E., Guenthner, C. J., Miyamichi, K., & Luo, L. (2014). Presynaptic partners of dorsal raphe serotonergic and GABAergic neurons. Neuron, 83(3), 645–662. doi:10.1016/j.neuron.2014.06.024

Yetnikoff, L., Cheng, A. Y., Lavezzi, H. N., Parsley, K. P., & Zahm, D. S. (2015). Sources of input to the rostromedial tegmental nucleus, ventral tegmental area, and lateral habenula compared: A study in rat. J Comp Neurol, 523(16), 2426–2456. doi:10.1002/cne.23797

Zahm, D. S., & Root, D. H. (2017). Review of the cytology and connections of the lateral habenula, an avatar of adaptive behaving. Pharmacol Biochem Behav, 162, 3–21. doi:10.1016/j.pbb.2017.06.004

Zahm, D. S., Schwartz, Z. M., Lavezzi, H. N., Yetnikoff, L., Parsley, K. P. J. B. S., & Function. (2014). Comparison of the locomotor-activating effects of bicuculline infusions into the preoptic area and ventral pallidum. 219(2), 511–526.

